# The Cerebellum and Striatum in Reward Processing: Caring About Being Right vs. Caring About Reward

**DOI:** 10.1101/2025.06.04.657253

**Authors:** Haroon Popal, Megan Quarmley, David V. Smith, Johanna Jarcho, Ingrid R. Olson

## Abstract

Emerging evidence indicates the cerebellum contributes to cognitive functions including social reward processing, yet its specific role relative to established reward regions like the ventral striatum remains undefined. We hypothesized the cerebellum would respond equivalently to both positive and negative social rewards. This prediction is grounded in classical findings that the cerebellum operates via supervised learning mechanisms that rely on error signals rather than traditional reward-based reinforcement. Using fMRI, we examined adolescents and young adults during a social prediction task where participants forecasted others’ opinions of them and received accuracy feedback. Findings reveal that both the ventral striatum and a subregion of the posterior cerebellum (Crus I and II) were sensitive to social rewards. However, unlike the ventral striatum, the cerebellum exhibited a more uniform response to feedback, treating correct predictions about being liked and disliked in a similar manner. No age-related differences were observed. These findings suggest the cerebellum processes social rewards distinctly from the ventral striatum, likely reflecting its computational emphasis on prediction errors rather than reward valence. This functional distinction advances our understanding of cerebellar contributions to social cognition and learning mechanisms.

## INTRODUCTION

Damage to the cerebellum early in life is associated with devasting social deficits (Olson et al., 2023) as well as an increased risk for autism spectrum disorder (ASD). Strikingly, neonatal cerebellum damage is the second largest risk factor for ASD, after having an identical twin with the disorder (Limperopoulos et al., 2007, 2014; Wang et al., 2014). Neuroimaging studies have implicated the cerebellum in a variety of tasks that capture how we understand others, including lower-order social cognitive processes such as biological motion perception (Jack & Pelphrey, 2015), and higher-order processes such as theory of mind (Metoki et al., 2022; Sokolov, 2018).

A region in the posterior lobe of the cerebellum, Crus I and II, has been strongly implicated in social mentalizing (Van Overwalle et al., 2015, 2019, 2020; Van Overwalle & Mariën, 2016). This region is functionally co-activated with socially-sensitive regions in the cerebrum, such as the temporoparietal junction (Heleven et al., 2021; Van Overwalle et al., 2020). Despite this evidence, the specific contribution of the cerebellum to social cognition – in other words, it’s computational role- is not known.

While the cerebellum is now known to be implicated in a variety of cognitive domains, the cytoarchitecture of the cerebellum is largely uniform (Ito, 2008), raising questions of how this singular structure of neurons can support functionally diverse processes. In the motor domain, the cerebellum’s cytoarchitecture has been shown to be well suited for performing supervised learning (Doya, 2000; Kawato et al., 1987; Raymond et al., 1996; Raymond & Medina, 2018; Wolpert et al., 1998). Briefly, granule cells in the cerebellar cortex receive inputs from the cerebrum and brainstem, and contexts are mapped onto specific optimal actions, building models of motor skills. Pukinje cells, the main output neurons in the cerebellum, send modulatory signals to the cerebrum, where action commands are created with the adjustments made by the cerebellum (Wolpert et al., 1998; Wolpert & Ghahramani, 2000). This cytoarchitecture allows the cerebellum to employ an internal forward model, which is used to predict the consequence of a planned action in order to quickly adjust signals to optimize such action. Some have speculated that because the uniform cytoarchitecture of the cerebellum is so well suited to performing supervised learning in this manner, it also does this for cognitive functions as well. This idea is termed *Universal Cerebellar Transform* (UCT) theory (Guell et al., 2018; Schmahmann, 2019). According to this theory, the cerebellum’s contribution to cognitive processes is to compute supervised learning algorithms that compare expected internal states to actual states, and then updating internal models (Ito, 2008). For example, relying on an internal forward model of social knowledge, one might adjust a joke they are about to tell co-workers to ensure that it is work-appropriate, thereby avoiding a social prediction error (e.g. feeling awkward), where one recognizes that in the current state (e.g. work) said joke would violate a social model (e.g. professional norms). An opposing theory to the UCT is the *Multiple Cerebellar Functionality* (MCF) theory, which proposes that a single type of cytoarchitecture can perform multiple computations, and therefore, serve different functions (Diedrichsen et al., 2019). An assumption of this theory is that the cerebellar cytoarchitecture can be biased towards performing alternate learning algorithms based on the input that a cerebellar region receives.

Until recently, there was little evidence that the cerebellum uses any algorithm besides supervised learning. However, recent work has hinted that the cerebellum is involved in reinforcement learning. Note that a major difference between supervised learning and reinforcement learning is a reward signal. In reinforcement learning, the receipt, or non-receipt of rewards act as a teaching signal (Niv, 2009). In supervised learning, an instructor that already knows the correct answers acts as the teaching signal. Traditionally, the striatum was thought to solely perform reinforcement learning via reward, while the cerebellum was thought to perform supervised learning without reward but that story in changing. Granule cells in lobule VI of the mouse cerebellum encode reward contexts including reward delivery, omission, and anticipation (Doya, 2000; Wagner et al., 2017); these signals were previously attributed to the striatum (Schultz, 2000). In addition, changes in D2 receptors in cerebellar Purkinje cells result in mice exhibiting abnormal social preferences (Cutando et al., 2022).

While reward is critical in reinforcement learning employed in the basal ganglia, it is unknown if reward is an important parameter for learning in the supervised learning employed in the cerebellum. Thus, a basic approach for understanding the cerebellum’s role in reinforcement learning is to ask: is the human cerebellum sensitive to reward?

### Current Study

In the current study we asked how the cerebellum responds to reward, and we contrasted this to how the ventral striatum responds to reward. Our rationale was that the ventral striatum has been established a key brain region in reinforcement learning and reward processing, hence it serves as a benchmark for such processing. If the cerebellum differs from the ventral striatum in its neural responses to reward, then the cerebellum is contributing something unique to reward processing.

Our fMRI study had two contrasting tasks: a social reward task and a non-social reward task (Quarmley et al., 2019). Traditionally, reward has been defined as a positive outcome for an individual, where they receive money or in the case of social cognition, feedback which makes the individual feel pleasant. Our task had a twist on this because you had to predict whether or not someone liked you, then you received feedback about the correctness of your prediction.

Discovering that someone does not like you is an unpleasant feeling, although if you were correct about them not liking you, it could provide a feeling of satisfaction – a type of internal reward -for being correct (Bhanji & Delgado, 2014), or it could provide you with information that is valuable in of itself (Shen et al., 2024; Tricomi & Fiez, 2012). By combining the two conditions of positive and negative valanced reward to create a valence-general correct condition, we can examine whether the cerebellum or the ventral striatum is more likely to be sensitive to this signal.

We asked a series of inter-related questions: (1) Does the cerebellum play a role in reward processing? We hypothesize that a reward-sensitive region would be found in Crus I and II, where social and cognition processes have previously been localized (Manto et al., 2024). (2) Does the cerebellum play a unique role in processing traditional and social rewards. To answer this question, representational similarity analysis (RSA) was used to test the hypothesis that the cerebellum follows a valence-general reward processing scheme, while the ventral striatum is particularly sensitive to positively valanced rewards. Because supervised learning operates without reward signals and instead relies on general error signals to adjust signals based on the magnitude of the error, we hypothesize that the cerebellum would treat positive and negative rewards more similarly than the ventral striatum. (3) Last, we asked whether there were any development differences in cerebellar processing of reward. A recent review has highlighted that cerebellar lesions earlier in life result in more dramatic social deficits than lesions that occur later life (Olson et al., 2023). This led to a cerebellar developmental cascade hypothesis which posits that cerebellar lesions earlier in life result in an unoptimized forward internal model, where the cerebellum is not able to build up internal social models. In later life, where internal models are already built, a deficit in cerebellar forward model processing would not result in deficits as the relevant social knowledge has already been learned. This study will examine if there are any differences in cerebellar processing of rewards between adolescents and young adults. Adolescence is a developmental time period in which social rewards are especially salient (Albert et al., 2013; Chein et al., 2011) and given evidence that the cerebellum is critical for prosocial behaviors in mice (Carta et al., 2019), we hypothesize that there would be differences between cerebellum activations for rewards between adolescents and young adults. Given that there is no literature on cerebellar processing of rewards in adolescence, this was an exploratory analysis.

## METHODS

### Participants

Participants were adolescents aged 11 - 16 (n = 37; females = 17; M age = 13.38) and young adults aged 18 - 36 (n = 45; females = 29; M age = 21.58) who were free of psychotropic medication use and had no contraindications for fMRI. Informed written parental consent and written participant assent were obtained from the adolescents, and informed written consent from the young adults prior to participation. All procedures were approved by the Institutional Review Board at Stony Brook University and were conducted in accordance with the Helsinki Declaration.

Since the cerebellum was not a region of interest at the time of data collection, some participants did not have the cerebellum in the fMRI field of view box. Eleven participants who did not have full cerebellum coverage were excluded. In addition, seven participants were excluded for having excessive motion in all of their fMRI runs, defined as maximum framewise-displacement greater than 4mm, and three participants were excluded due to incomplete fMRI data runs and fMRI artifacts. This resulted in a final sample of adolescents (n = 32; females = 15; M age = 13.38) and young adults (n = 29; females = 21; M age = 21.21; SD).

### Procedure

Prior to the experimental session, participants were told they were completing a social evaluation study and were asked to submit a digital picture of themselves that would be sent to other purported participants their age across the country. Participants believed that these peers would receive a text message asking them to view the photo and indicate whether they thought they would “like” or “dislike” the participant. The picture would then disappear after five minutes. At the beginning of the experiment session, participants were told that they would be asked to guess which peers “liked” or “disliked” them and that they would also be completing a monetary guessing task. Participants underwent mock scanning to gain familiarity with the MRI environment. Participants then underwent fMRI while completing the monetary and social reward tasks in a counterbalanced order. At the end of the session, participants responded to questions about their experience with the task to ensure they were engaged and believed the credibility of the peer feedback. Nearly all (85%) participants had high levels of task engagement and believed they were receiving feedback from actual peers. Participants were then debriefed.

#### fMRI-Based Monetary and Social Reward Tasks

The monetary and social reward tasks (Fig. 1) were administered using Eprime software (E-Prime 2.0). There were four conditions (monetary positive, monetary negative, social positive, and social negative) that were presented in a counterbalanced order. Each condition included 30 trials. Each task was completed across two, 4.55-min runs. Each run included two blocks: one block of monetary positive or social positive trials, and one block of monetary negative or social negative trials (15 trials per block). Trials were separated by a variable duration intertrial interval (1,100–11,600 ms; *M* = 3,500 ms).

**Figure 1.**
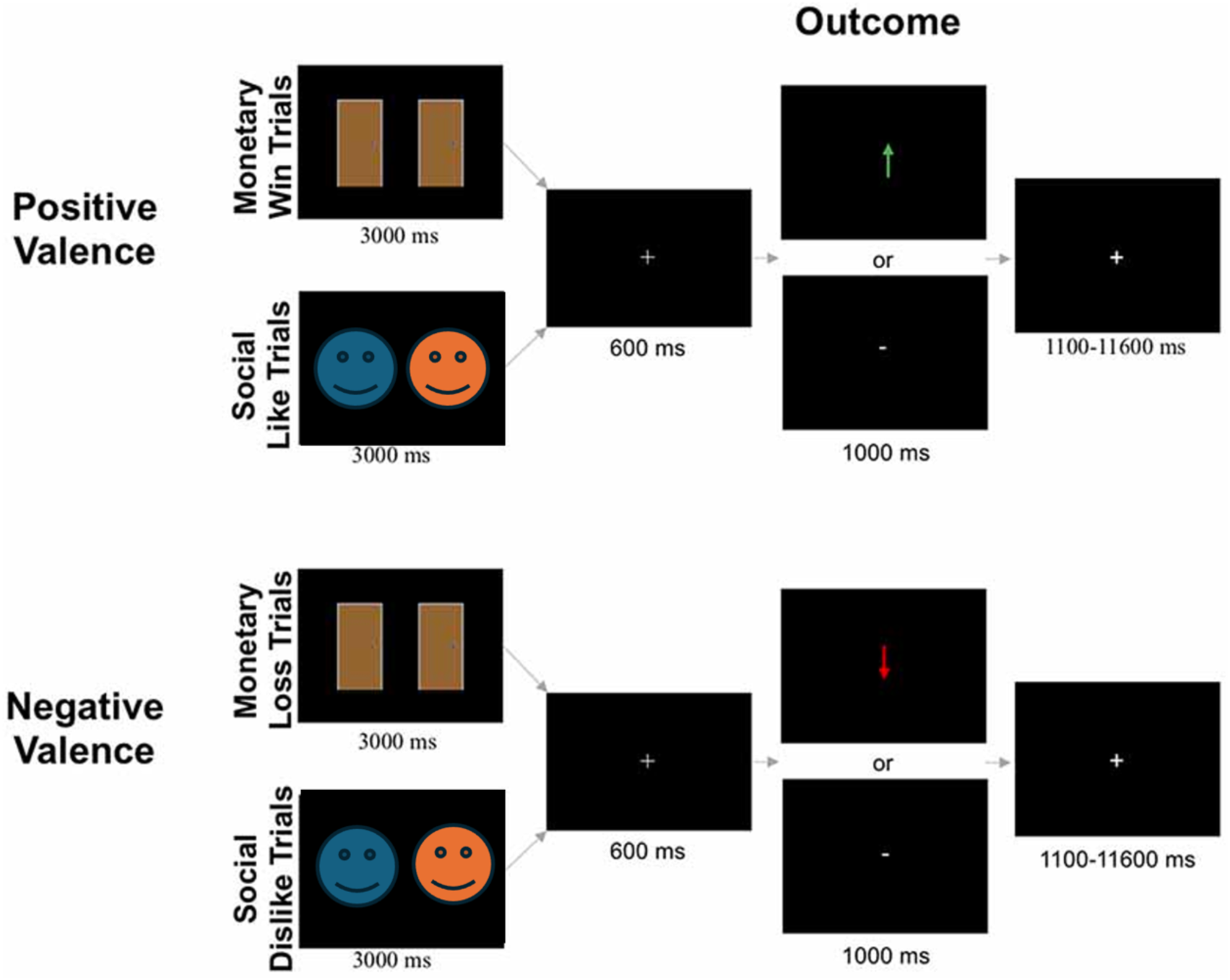
The monetary and social doors task. Positive valanced trials required participants to select which door/face resulted in a positive reward (i.e. winning money or being liked). Negative valanced trials required participants to select which door/face resulted in a negative reward (i.e. losing money or being disliked). In the outcome phase of the trial, an arrow (green or red for positive or negative valence, respectively) indicated a correct outcome, and “-” indicated an incorrect outcome. Due to bioRxiv’s policy on avoiding the inclusion of photographs of people in figures, a blue and orange slimy face is shown here. In the actual task, photographs of real people from the National Institute of Mental Health’s Child Emotional Faces picture set (Egger et al., 2011 and internet databases of non-copyrighted images were used.

#### Monetary Reward Task

At the beginning of each block, participants were informed if the block contained monetary positive trials or monetary negative trials. In the positive valanced monetary blocks, participants were instructed to choose the door behind which there was a $0.25 prize. In the negative valanced monetary blocks, participants were instructed to choose the door behind which there was a $0.25 monetary loss. Each trial began with the presentation of two identical doors (3,000 ms). Participants then used a button box to select either the left or right door on the screen. After stimulus offset, a fixation cross was presented for 600 ms before participants received feedback about the accuracy of their choice (1,000 ms). Participants were told that there were three possible outcomes for each monetary win/loss trial: (1) both doors contained a $0.25 monetary win/loss; (2) one door contained a $0.25 monetary win/loss while the other door resulted in a break-even outcome (i.e., neither win nor loss); or (3) both doors resulted in a break-even outcome. This ensured that the feedback the participant received would only be informative about the door they chose and not the door they *did not* choose. For example, if a participant chose a door and received feedback indicating a break-even result, the other door could have been a win/loss door (consistent with trial scenario above) or it could have been another break-even door (consistent with trial scenario above). In monetary positive trials, feedback was either a green arrow pointing upward (↑) meaning the participant correctly selected the monetary win door, or a white horizontal dash (-), which indicated incorrect selection of the break-even door, resulting in no monetary win. In monetary negative trials, correct selection of the monetary loss door was indicated by a red arrow pointing downward (↓), while incorrect selection of the break-even door resulting in no monetary loss was indicated by a white horizontal dash (-).

#### Social Reward Task

The social positive and negative conditions were identical to the monetary positive and negative conditions, respectively, except pictures of gender-matched peers (i.e., two female faces or two male faces) were presented instead of doors. The social reward task consisted of 120 images of age-matched peers compiled from multiple sources [National Institute of Mental Health’s Child Emotional Faces picture set (Egger et al., 2011) and internet databases of non-copyrighted images]. The pictures of purported peers had positive facial expressions, were cropped so that individuals were pictured from their shoulders up, and were edited to have an identical solid gray background. Smiling faces were used because they are common in social reward tasks (Distefano et al., 2018; Jarcho et al., 2015; Richards et al., 2013) and are subject to less misinterpretation than neutral faces (Davis et al., 2016; Rapee & Heimberg, 1997). Images were constrained to a standard size (2.75 inch width × 4 inch height). There were an equal number of trials with male and female peers across the social positive and negative conditions (30 pairs each, 60 total).

At the beginning of each block of trials, participants were informed if the block contained positive valanced social like trials or negative valanced social dislike trials. In social positive blocks, participants were instructed to choose the peer that liked them. In social negative blocks, they were instructed to choose the peer that disliked them. Participants were told that there were three possible outcomes for each trial: (1) both people said they would like/dislike the participant; (2) one person said they would like/dislike the participant while the other person never rated the participant; or (3) neither person rated the participant. In social positive trials, correct selection of the person who said they would like the participant was indicated by a green arrow pointing upward (↑). In social negative trials, correct selection of the person who said they would dislike the participant was indicated by a red arrow pointing downward (↓). In both social positive and negative conditions, incorrect selection of the person who never rated the participant was indicated by a white horizontal dash (-).

### MRI Acquisition

Functional images were acquired using a 3T Siemens PRISMA MRI scanner. Blood Oxygenation Level-Dependent (BOLD) sensitive functional images were acquired using a gradient echo-planar imaging (EPI) sequence (224 mm in FOV, TR = 2,100 ms, TE = 23 ms, voxel size of 2.3 × 2.3 × 3.5 mm^3^, flip angle = 83°, interleaved slice acquisition). Each run included 37 functional volumes. To facilitate anatomical localization and coregistration of functional data, a high-resolution structural scan was acquired (sagittal plane) with a T1- weighted magnetization-prepared rapid acquisition gradient echo (MPRAGE) sequence (250 mm in FOV, TR = 1,900 ms, TE = 2.53 ms, voxel size of 1.0 × 1.0 × 1.0 mm^3^, flip angle = 9°).

### MRI Preprocessing

Structural and functional MRI preprocessing was completed with fMRIPrep (Esteban et al., 2019). Complete information on preprocessing can be found in the supplementary materials. Briefly, structural images were normalized to the MNI152NLin2009cAsym template. Functional images were slice-time corrected, and co-registered and resampled to their original, native space. No smoothing was applied in the preprocessing nor the analysis stages. Functional runs with greater than 4 mms of maximum framewise displacement were excluded. All functional runs were examined to ensure full brain coverage, including the cerebellum.

### Data Analysis

#### Regions of Interests

Regions of interests (ROIs) were defined from three separate sources since there were three groups of brain regions that would be relevant for this study (Fig. 2). The striatum was a region of interest because of its well-known involvement in reward processing. Striatum ROIs were defined using the striatum atlas from FSL (Tziortzi et al., 2011), which parcellates the striatum in ventral, anterior dorsal, and posterior dorsal subregions. All three subregions from this atlas were included as separate ROIs, with left and right regions being combined to create bilateral ROIs. The ventromedial prefrontal cortex (vmPFC) is also relevant for reward processing and is thought to process the value of rewards (Smith et al., 2010, 2014). Therefore, an ROI was created using the “reward” term in Neurosynth to create a brain map (Yarkoni et al., 2011). From this whole brain map, subregions were created using the “expand_mask” command from nltools (https://nltools.org/), and the right vmPFC was extracted. The “social” term was used to create a brain map in Neurosynth for ROIs which would be relevant for social cognition. Left and right homologs were combined to create bilateral ROIs. This resulted in ROIs for the bilateral temporoparietal junction (TPJ), bilateral anterior temporal lobes (ATL), bilateral posterior cingulate cortex (PCC), dorsal medial prefrontal cortex (dmPFC), and the right amygdala. Lastly, the multi-domain task battery (MDTB) functional atlas (King et al., 2019) was used to create 10 parcellations for the cerebellum. Please note that it is difficult to use Neurosynth for cerebellar ROIs, given that many fMRI studies fail to include the cerebellum in the FOV. Th MDTB atlas is a task-based parcellation of the cerebellum based on 47 task conditions. Of note, this cerebellum atlas has specific regions that have been labeled as either motor or specific cognitive regions. Two motor region ROIs (region 1 and 2 from the MDTB atlas) were selected to act as control regions. Regions 5-10 from the MDTB atlas, which overlap with lobule VI, Crus I, and Crus 2, were selected as ROIs because these regions have previously been shown to be sensitive to reward and/or social cognition (Carta et al., 2019; Van Overwalle et al., 2015; Wagner et al., 2017). In total, 17 ROIs were created to capture reward, social, and cerebellum regions that may be relevant to this study.

**Figure 2.**
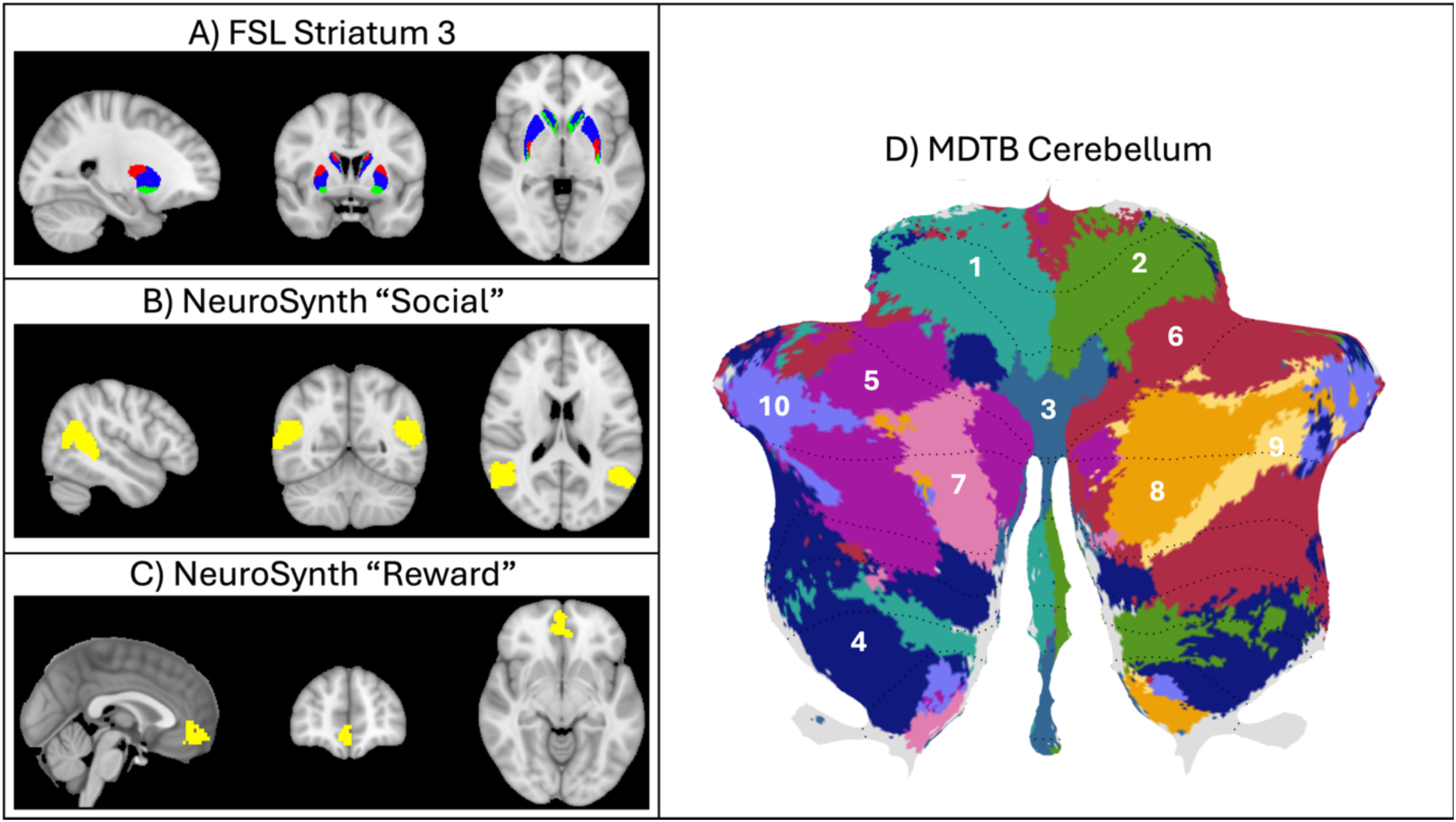
Regions of interest. (A) Striatum 3-part parcellation from FSL atlas, showing subregions for ventral (green), anterior dorsal (blue), and posterior dorsal (red) striatum. (B) NeuroSynth “social” term with TPJ extracted. (C) NeuroSynth “reward” term with vmPFC extracted. (D) Multi-domain task based parcellation atlas of the cerebellum (used with permission from King et al., 2019).

#### First Level Analyses

First-level analyses were done using a general linear model (GLM) to examine the overall response for each condition of interest using the python based nilearn package. GLMs were defined with the “slice_time_ref” parameter set to “0.5”, to account for the slice time correction method used with fmriprep. For each task (monetary and social) and for each valence condition (positive and negative), the regressors of interest were the decision phase of each trial and the outcome phase of each trial (correct and incorrect), resulting in six regressors: positive decision, positive correct outcome, positive incorrect outcome, negative decision, negative correct outcome, and negative incorrect outcome. Along with these six regressors, nuisance regressors were included in the design matrix for each run which included a constant, a linear regressor, and six motion regressors for movement and rotation along the x, y, and z axes.

Design matrices and runs for each task were concatenated. All regressors were convolved with the canonical hemodynamic response function. Whole brain z-statistic maps were created for each regressor of interest, indicating whether each voxel was significantly greater than zero.

Contrast maps were also created for positive valanced correct vs incorrect trials, and for all (positive and negative) correct vs incorrect trials, for each task.

#### Cerebellum Spatial Normalization

The structural architecture of the cerebellum is highly unique with its highly foliated gyri. Given that most preprocessing software was developed for the cerebrum, special attention must be paid to cerebellar normalization. We used the spatially unbiased infratentorial template (SUIT) SPM 12 toolbox (Diedrichsen, 2006) to normalize the cerebellum based on each participant’s T1 scan. Afterwards, each first-level statistical map was resliced into the SUIT cerebellum space, and cerebellar analyses were conducted in this new space.

#### Univariate Analysis

To examine whether an ROI was significantly more responsive for positive correct over incorrect trials and for all correct over incorrect trials, univariate contrasts were conducted as a first-level analysis for each participant. Then, a two-way ANOVA was completed to examine if there was a significant difference between all the ROIs, tasks, or an interaction between the two variables. Post-hoc one-sample t-tests were done to see if each ROI’s contrast activation was significantly greater than zero, and paired t-tests were done to examine task differences between each ROI. False discovery rate (FDR) correction was used to account for multiple comparisons for each contrast, using a family-wise error rate of *p* < .05 for the post-hoc tests. An additional set of ANOVAs were completed similar to the ANOVA described above, but examined main effects for ROI, group (adolescent vs young adult), and their interaction, independently for each task.

#### Representational Similarity Analysis

Representational similarity analysis (RSA) was used to examine whether the cerebellum and striatum process information about reward similarly. All first-level analyses were conducted using the nilearn python toolbox. To test the hypothesis that the cerebellum is a valence-neutral error processor, we compared multi-voxel pattern responses between positive and negative correct trials. Activity from voxels within an ROI were extracted from t-statistic maps for positive and negative correct trials in every run and task. The positive and negative condition responses were correlated to each other, using Spearman’s rho, for a given ROI, across runs, but not within the same run. For example, the multi-voxel pattern response for positive correct trials in Run 1 of the social task was correlated with the response for negative correct trials in Run 2 of the social task. It has been suggested that correlating conditions between runs in this manner is able to address the autocorrelations present within runs for multi-variate pattern analyses, especially in a block design task where conditions are not explicitly random (Dimsdale-Zucker & Ranganath, 2019). Complementary comparisons were then averaged (e.g. Run 1 positive correct to Run 2 negative correct and Run 2 positive correct to Run 1 negative correct).

Second level RSA was done in R with similar methods to the univariate analysis. The positive and negative reward condition correlations for each participant were inputted into a two-way ANOVA, examining main effects for ROI, task, and an interaction between the two variables. Post-hoc one-sample t-tests were done to see if each ROI’s positive to negative reward correlation was significantly greater than zero, and paired t-tests were done to examine task differences between each ROI. An FDR correction was used to account for multiple comparisons for each contrast and task, using a family-wise error rate of *p* < .05 for the post-hoc tests.

## RESULTS

### Age Differences in Cerebellar Processing

While examining differences in cerebellar activity between adolescents and young adults was an exploratory aim, no group differences were observed in the univariate analyses.

Therefore, for the rest of the analyses, the adolescent and young adult samples were combined and analyzed together. Results for the group analysis are available in the supplementary materials.

### Sensitivity to Monetary and Social Reward

For the positive correct > positive incorrect contrast, the overall ANOVA showed significant main effects for task (F(1, 2040) = 9.30, *p* = .002; η^2^ (partial) = 4.54e-03), and ROIs (F(16, 2040) = 2.63, p < .001; η^2^ (partial) = 0.02), but not for the interaction (F(16, 204) = 0.46, p = .965; η^2^ (partial) = 3.61e-03). In the post-hoc analyses, there was no significant differences

in responses for the positive correct > positive incorrect contrast between the monetary and social tasks for any of the individual ROIs, after FDR correction. ROIs that were significantly more activated for positive correct over positive incorrect trials in the monetary task included the right amygdala, vmPFC, CB region 8, CB region 9, ventral striatum, and the anterior dorsal striatum. All the ROIs were significantly more activated for positive correct over positive incorrect trials in the social task, except for the ATL (Fig. 3A).

**Figure 3.**
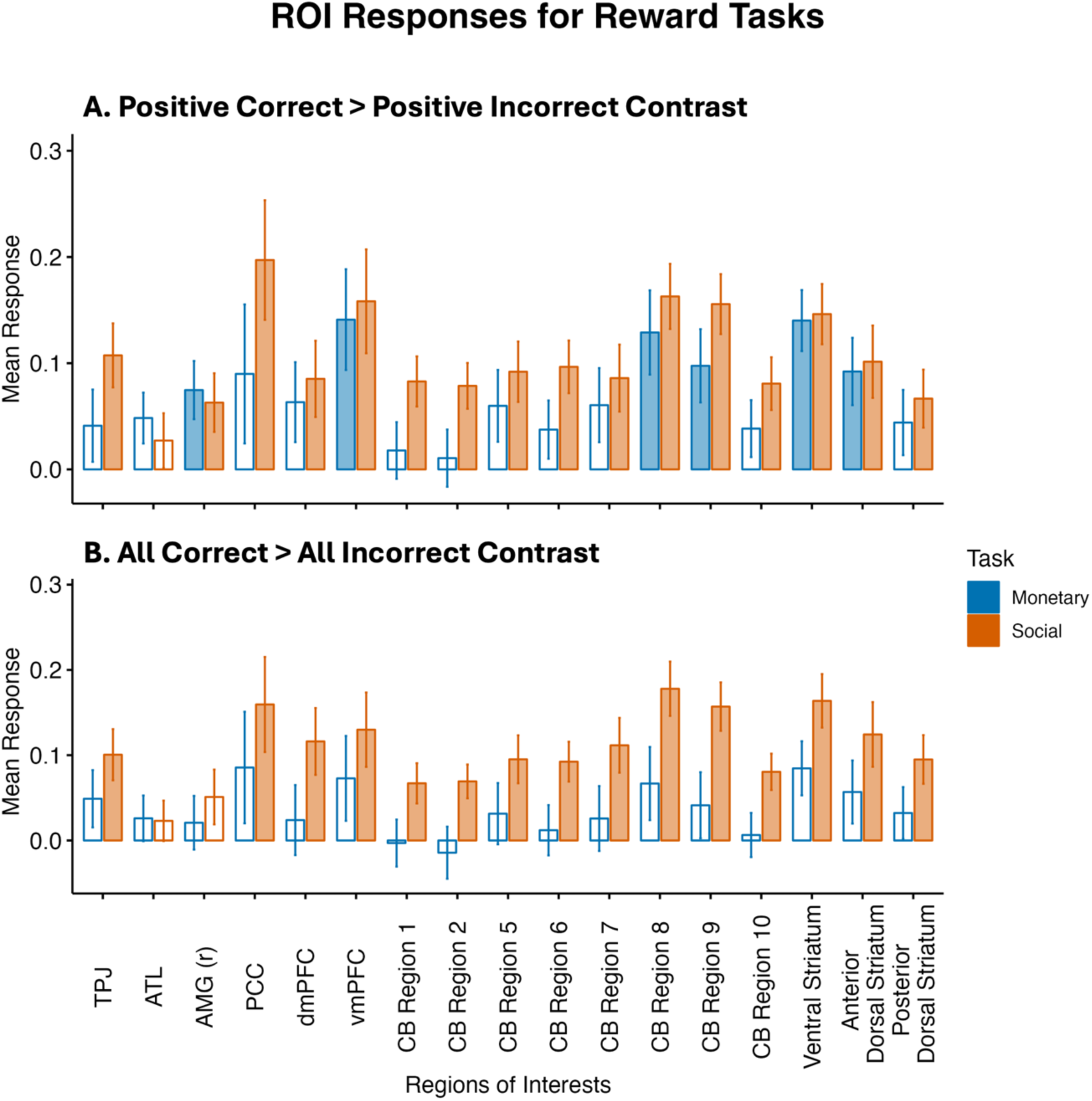
Univariate results. (A) positive correct > positive incorrect; and (B) all (positive and negative) correct > all (positive and negative) incorrect contrasts for the monetary (blue) and social (orange) tasks. Filled in bars indicate significant one-sample t-tests for the specific ROI and task, *p* < .05, FDR corrected. TPJ = bilateral temporoparietal junction, ATL = bilateral anterior temporal lobes, AMG (r) = right amgydala, vmPFC = ventromedial prefrontal cortex, PCC = posterior cingulate cortex, CB = cerebellum.

For the all correct > all incorrect contrast of the monetary reward task, the overall ANOVA showed significant main effects for group (F(1, 2040) = 34.28, *p* < .001; η^2^ (partial) = 0.02), and ROIs (F(16, 2040) = 1.86, *p* = .02; η^2^ (partial) = 0.01), but not for the interaction (F(16, 2040) = 0.32, *p* = .995; η^2^ (partial) = 2.48e-03). In the post-hoc analyses, there were no significant differences between tasks for any of the ROIs. No ROIs were significantly more activated for all correct versus all incorrect trials in the monetary task. In the social task, all ROIs except for the ATL and right amygdala were significantly more activated for all correct than all incorrect trials (Fig. 3B).

### Cerebellum vs Striatum Reward Processing

Across all the tasks and contrasts, two regions were most consistently sensitive to correct trials: CB region 8 and the ventral striatum. To differentiate these regions, RSA was used by comparing how similarly each ROI responds to positive and negative correct trials. CB region 2 was also used as a control ROI, as it has been well-established to be involved in motor functions (King et al., 2019). A two-way ANOVA was used to also examine differences between ROIs and tasks, and for a possible ROI by task interaction. The overall ANOVA showed significant main effects for ROI (F(2, 360) = 22.66, *p* < .001; η^2^ (partial) = 0.11) and task (F(1, 360) = 7.30, *p* = 0.007; η^2^ (partial) = 0.02), and a significant interaction (F(2, 360) = 4.04, *p* = 0.018; η^2^ (partial) = 0.02). Taken together, these findings suggest that the ROIs behave differently from each other, and between tasks. To examine which ROI had the greater positive and negative reward similarity, and a difference between monetary and social tasks, a direct comparison between ROIs was done.

Across both tasks, CB region 8, CB region 2, and the ventral striatum all had significantly similar activation patterns between positive and negative correct trials (Fig. 4). This similarity was significantly higher in CB region 8 (M = 0.09, SD = 0.12) compared to CB region 2 (M = 0.07, SD = 0.06) and the ventral striatum (M = 0.02, SD = 0.05). Lastly, within CB region 8, the similarity between social positive correct and social negative correct trials was greater than monetary positive correct and monetary negative correct trials (t(60) = 2.56, *p* = .0128). Overall, this indicates that between these three ROIs, CB region 8 exhibits the strongest profile of a valence-general error processor that treats positive valanced feedback and negative valanced feedback in a similar manner, while being more responsive to the magnitude of the feedback.

**Figure 4.**
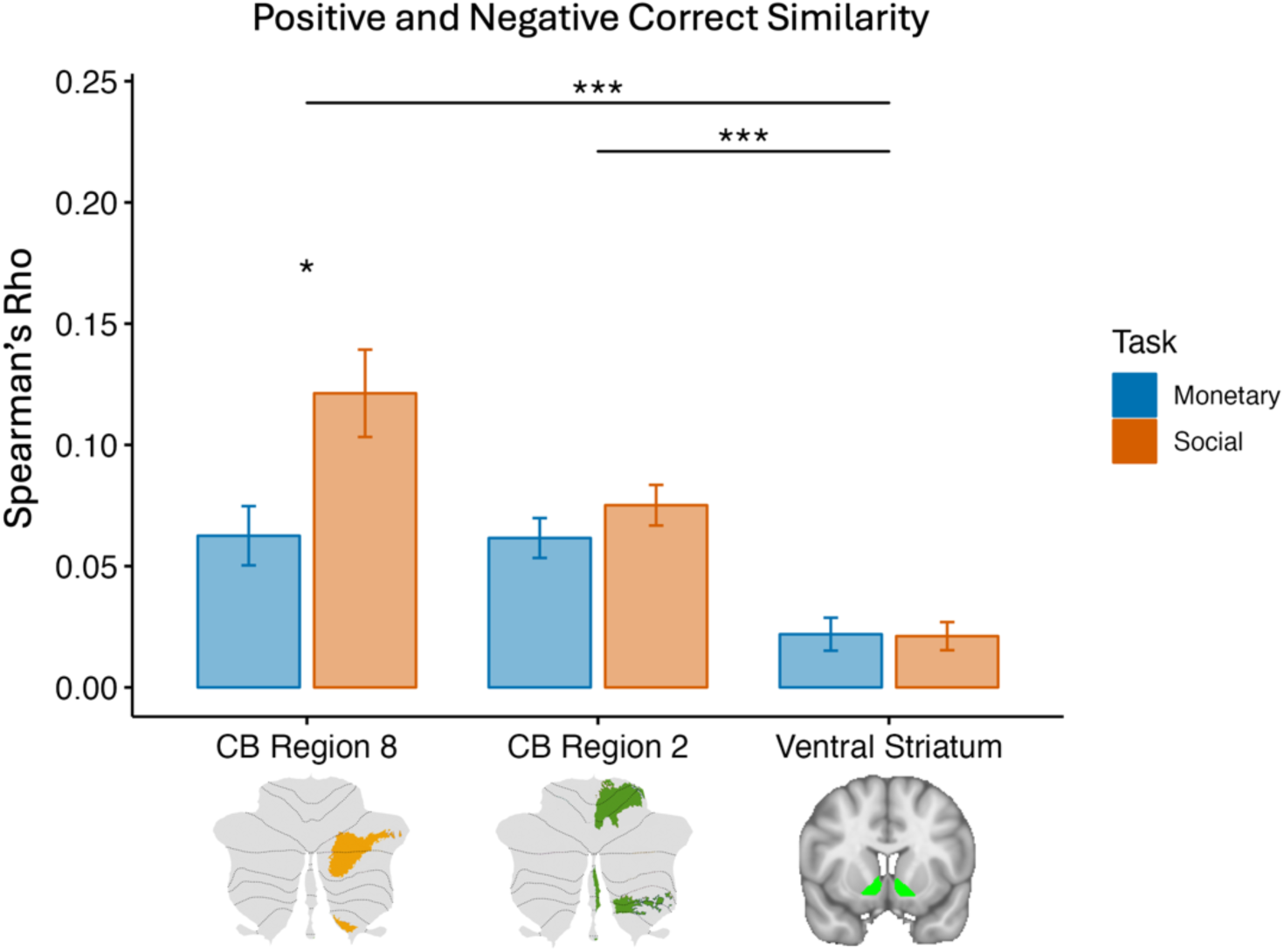
The similarity between positive and negative correct trials, in three regions. Cerebellar region 8 and the ventral striatum were most sensitive to correct outcomes (CB region 2 is a motor region that was used as a control) and were used as the basis of our comparison. We used RSA to compare positive correct to negative correct trials in the monetary (blue) and social (orange) tasks. The overall ANOVA revealed significant main effects for ROI and task, and a significant interaction. CB Region 8 had significantly greater similarity between responses for positive correct and negative correct trials, than CB region 2 and the ventral striatum. This evidence supports the universal cerebellar transform theory of a valence-general supervised learning processor, as it differentiates from the more valence-sensitive ventral striatum which has been implicated in reinforcement learning. The similarity between positive and negative correct trials was greater in the social task than the monetary task for CB region 8. Filled in bars indicate a significant one-sample t-test indicating greater than 0 similarity, *p* < .05 adjusting for multiple comparisons. Asterisks indicate significant paired t-tests between ROIs or tasks. Error bars represent standard error. * = *p* < .05, ** = *p* < .01, *** = *p* < .005, adjusting for multiple comparisons. CB = cerebellum.

## DISCUSSION

In this study we tested adolescents and young adults on a social reward task. We asked whether portions of the cerebellum are sensitive to social rewards and if so, how do these cerebellar regions use reward in a way that is unique as compared to the basal ganglia’s processing of reward? Neuroanatomical models of reward processing have focused heavily on the striatum, part of the basal ganglia, and medial regions of prefrontal cortex (Dayan, 2009; Graybiel & Grafton, 2015; Peak et al., 2019; Wise, 2004). However, there is emerging work showing that the cerebellum plays a role in reward processing and reinforcement learning (Carta et al., 2019; Kostadinov et al., 2019; Wagner et al., 2017). First, we asked does the cerebellum play a role in reward processing? We found that portions of the posterior lobe of the cerebellum, namely Crus I and Crus II, are sensitive to social rewards (Fig. 3). The location of this effect is consistent with prior work showing that Crus I and II are involved in higher level cognition, including theory of mind and language (King et al., 2019; Van Overwalle et al., 2015).

Second, we asked, does the cerebellum play a unique role in processing monetary and social rewards? RSA results suggest that Crus I/II regions are more sensitive to *domain general* rewards than the striatum. That is, cerebellar Crus I/II are more concerned with the “correct” answer, rather than correct answers that result in positive outcomes, suggesting that the cerebellum does indeed use rewards in a way that is distinct from the striatum. This is interesting because it re-casts the role of rewards into a teaching signal, used to refine behavior, rather than as a signal used to motivate behavior. This also provides modest evidence for the cerebellar uniform transform theory with the cerebellum employing a supervised learning algorithm minimizing error, as it distinguishes the reward signals observed in cerebellum from those seen in the ventral striatum which employs reinforcement learning for positive reward optimization.

The cerebellum and the striatum are strongly interconnected with a loop-like architecture: the dentate nucleus of the cerebellum sends dense disynaptic projections to the striatum, while a different region of the basal ganglia, the subthalamic nucleus, sends dense disynaptic projections to cerebellar cortex (Bostan et al., 2010; Bostan & Strick, 2018). This cerebellar-basal ganglion network is topographically organized such that motor, cognitive, and affective territories of each node in the network are interconnected (Bostan et al., 2010; Bostan & Strick, 2018), providing evidence that evolution precisely aligned these brain regions to work together.

Last, we asked whether there were any developmental differences. We found no difference between adolescents and adults. Perhaps by adolescence, the cerebellum has refined its joint function with the striatum and its sensitivity to rewards is identical to an adult’s. We predict that age differences will be more apparent in younger age groups, given the explosion of growth seen before the age of 5, with continued but slower development until around the age of 13 (Sathyanesan et al., 2019). We do not know the developmental timetable of when the cerebellum is functional differentiated (Supekar et al., 2009). In addition, age effects on reinforcement learning correlates have been shown to be nonlinear between the ages of 9-18 (Rosenblau et al., 2018), suggesting that the adolescent sample studied in this project may be at the bottom of a “U” shaped trajectory of development.

The apparent contradiction between the cerebellum’s established role in novel skill acquisition and its reliance on supervised learning mechanisms raises fundamental questions about cerebellar function in reward processing. The supervised learning framework presupposes an instructive signal indicating optimal performance parameters—yet how does this mechanism operate when acquiring entirely new social or cognitive skills where the “correct” execution remains undefined? A particularly compelling resolution emerges in the form of the “super learning” hypothesis (Caligiore et al., 2019), which postulates that the striatum and cerebellum function as complementary learning systems with strategic interactions during early acquisition phases. Within this framework, the striatum—operating via reinforcement learning principles that balance exploration and exploitation—transmits reward signals to the cerebellum, effectively designating successful actions as “correct” and thereby serving as the supervisor necessary for cerebellar supervised learning. This interactive neural architecture creates a sophisticated learning system wherein the basal ganglia facilitates exploratory behavior while the cerebellum concurrently establishes more stable response patterns. The differential learning rates between these systems—with the cerebellum having a slower learning rate than the striatum (Taylor & Ivry, 2014)—may represent an evolutionary optimization allowing the cerebellum to maintain representations of consistently successful actions while the basal ganglia continues to explore alternative response possibilities. This theoretical model elegantly accounts for the structural and functional connectivity observed between these neural structures and provides a neurobiologically plausible explanation for their complementary contributions to reward-based learning.

### Limitations and Suggestions for Future Research

First, our reward task did not have an explicit learning component, which would have allowed participants to choose conditions which would have allowed them to optimize rewards. A task with some patterned repetition that allows participants to build models based on their performance would have been better situated to answer the research question of how the cerebellum contributes uniquely to learning compared to the striatum. Two-stage reinforcement learning tasks offer a possible avenue of differentiating the cerebellum as these tasks can be used to separately pinpoint signals related to model-free and model-based learning. Given the cerebellum’s role in supervised learning, we hypothesize that the cerebellum would mimic model-based learning as seen in the prefrontal cortex more closely than model-free learning (Doll et al., 2015; Gläscher et al., 2010). Second, the monetary task was less visually interesting than the social task. The door stimuli were visually identical, while the faces were all unique.

Participants may have selected doors at random, whereas in the social condition, they had more features to rely on for making a decision. This may explain some of the observed stronger effects in the social task than the monetary task, and presents a limitation in making claims about the social specificity of our findings in terms of social versus non-social reward processing. A follow-up study by the original authors of this study is using unique doors (Wyngaarden et al., 2024).

### Future Directions

This study is novel in that it has shown that the cerebellum is directly involved in reward processing, in a way that is unique from the striatum. A simple next step would be to examine cerebellar processing during social reinforcement learning, to examine if there are differences in how the cerebellum and ventral striatum contribute to learning. Future research should be aimed at developing tasks that can directly differentiate cerebellar from striatal processes. More naturalistic learning paradigms should be adopted in tasks as well. Social situations are incredibly dynamic. This study took a more basic approach in order to pinpoint a very fundamental signal of reward. Naturalistic experiments can offer the opportunity to capture more dynamic reward signals. This study has shown that being correct about a negative outcome is still rewarding. There are likely other aspects of social situations that can likewise be rewarding because they are social, and which do not exist in other non-social contexts.

A direct examination of the cerebellum’s contribution to social learning is also warranted, as several contributions are possible. This contribution could be to more lower-level social cognition, by aiding in the processing of biological motion (Jack et al., 2017; Jack & Pelphrey, 2015), a necessary component of visual social learning. It could also be more higher-level by helping to build up social knowledge, as we have shown that the cerebellum is reliably activated when predicting appropriate social interactions for individuals of a specific social relationship.

The cerebellum has already been implicated in social learning (Rosenblau et al., 2018), but a direct examination of its contribution has yet to be studied. Greater white matter connections, measured via diffusion weighted imaging, between medial portions of the posterior cerebellum and the ventral tegmental area have been positively correlated with anxiety and depression (Hoffman et al., 2024), suggesting that medial portions of the posterior cerebellum play a role in socioaffective regulation via reward networks.

Some of the earliest work in understanding the neural correlates of ASD implicated the cerebellum (Allen & Courchesne, 2003; Courchesne & Allen, 1997). As social learning challenges are common in ASD (Kruppa et al., 2019; Lin et al., 2012), the cerebellum’s role should also be considered. The cerebellum could be unique in its implications in ASD, compared to other brain regions, as the cerebellum’s role in supervised learning could be involved in multiple ASD symptoms. Sensory challenges may be related to differences in the cerebellum, of modulating sensory prediction errors, while difficulties in maintaining active conversations could come from prediction errors relating to reading body language, and making inappropriate comments could be from difficulties in building up mental models of social norms.

## Conclusion

The goal of this study was to distinguish the cerebellum’s role in reward processing from that of the basal ganglia. We found that subregions of the posterior cerebellum are sensitive to reward outcomes, and that these regions are particularly sensitive to correct outcomes, regardless of whether the outcomes are positive or negative. This is distinct from the striatum, which showed a weaker similarity between positive and negative rewards. Overall, this supports the universal cerebellar transform theory and shows that the cerebellum acts as a general error detector, more in line with supervised learning, than reinforcement learning.

## Acknowledgements

We would like to thank Yin Wang, Allison Jack, and Jason Chein for their feedback in reviewing the manuscript and analysis suggestions.

## References

Albert, D., Chein, J., & Steinberg, L. (2013). The Teenage Brain: Peer Influences on Adolescent Decision Making. Current Directions in Psychological Science, 22(2), 114–120. 10.1177/0963721412471347

Allen, G., & Courchesne, E. (2003). Differential effects of developmental cerebellar abnormality on cognitive and motor functions in the cerebellum: An fMRI study of autism. American Journal of Psychiatry, 160(2), 262–273. 10.1176/appi.ajp.160.2.262

Bhanji, J. P., & Delgado, M. R. (2014). The social brain and reward: Social information processing in the human striatum. WIREs Cognitive Science, 5(1), 61–73. 10.1002/wcs.1266

Bostan, A. C., Dum, R. P., & Strick, P. L. (2010). The basal ganglia communicate with the cerebellum. Proceedings of the National Academy of Sciences of the United States of America, 107(18), 8452–8456. 10.1073/pnas.1000496107

Bostan, A. C., & Strick, P. L. (2018). The basal ganglia and the cerebellum: Nodes in an integrated network. Nature Reviews Neuroscience, 19(6), 338–350. 10.1038/s41583-018-0002-7

Carta, I., Chen, C. H., AmandaL.Schott, Dorizan, S., & Khodakhah, K. (2019). Cerebellar modulation of the reward circuitry and social behavior. Science, 363(January). 10.1126/science.aav0581

Chein, J., Albert, D., O’Brien, L., Uckert, K., & Steinberg, L. (2011). Peers increase adolescent risk taking by enhancing activity in the brain’s reward circuitry. Developmental Science, 14(2), F1–F10. 10.1111/j.1467-7687.2010.01035.x

Courchesne, E., & Allen, G. (1997). Prediction and preparation, fundamental functions of the cerebellum. Learning & Memory (Cold Spring Harbor, N.Y.), 4, 1–35. 10.1101/lm.4.1.1

Cutando, L., Puighermanal, E., Castell, L., Tarot, P., Belle, M., Bertaso, F., Arango-Lievano, M., Ango, F., Rubinstein, M., Quintana, A., Chédotal, A., Mameli, M., & Valjent, E. (2022). Cerebellar dopamine D2 receptors regulate social behaviors. Nature Neuroscience, 25(7), 900–911. 10.1038/s41593-022-01092-8

Davis, F. C., Neta, M., Kim, M. J., Moran, J. M., & Whalen, P. J. (2016). Interpreting ambiguous social cues in unpredictable contexts. Social Cognitive and Affective Neuroscience, 11(5), 775–782. 10.1093/scan/nsw003

Dayan, P. (2009). Dopamine, reinforcement learning, and addiction. Pharmacopsychiatry, 42 *Suppl 1*, 56–65. 10.1055/s-0028-1124107

Diedrichsen, J. (2006). A spatially unbiased atlas template of the human cerebellum. NeuroImage, 33(1), 127–138. 10.1016/j.neuroimage.2006.05.056

Diedrichsen, J., King, M., Hernandez-castillo, C., Sereno, M., & Ivry, R. B. (2019). Universal transform or multiple functionality? Understanding the contribution of the human cerebellum across task domains. Neuron, 102(5), 918–928. 10.1016/j.neuron.2019.04.021

Dimsdale-Zucker, H. R., & Ranganath, C. (2019). Representational similarity analyses: A practical guide for functional MRI applications. In Handbook of In Vivo Plasticity Techniques (pp. 509–525). Elsevier B.V. 10.1016/b978-0-12-812028-6.00027-6

Distefano, A., Jackson, F., Levinson, A. R., Infantolino, Z. P., Jarcho, J. M., & Nelson, B. D. (2018). A comparison of the electrocortical response to monetary and social reward. Social Cognitive and Affective Neuroscience, 13(3), 247–255. 10.1093/scan/nsy006

Doll, B. B., Duncan, K. D., Simon, D. A., Shohamy, D., & Daw, N. D. (2015). Model-based choices involve prospective neural activity. Nature Neuroscience, 18(5), 767–772. 10.1038/nn.3981

Doya, K. (2000). Complementary roles of basal ganglia and cerebellum in learning and motor control. Current Opinion in Neurobiology, 732–739. 10.1016/S0959-4388(00)00153-7

Egger, H. L., Pine, D. S., Nelson, E., Leibenluft, E., Ernst, M., Towbin, K. E., & Angold, A. (2011). The NIMH Child Emotional Faces Picture Set (NIMH-ChEFS): A new set of children’s facial emotion stimuli. International Journal of Methods in Psychiatric Research, 20(3), 145–156. 10.1002/mpr.343

Esteban, O., Markiewicz, C. J., Blair, R. W., Moodie, C. A., Isik, A. I., Erramuzpe, A., Kent, J. D., Goncalves, M., DuPre, E., Snyder, M., Oya, H., Ghosh, S. S., Wright, J., Durnez, J., Poldrack, R. A., & Gorgolewski, K. J. (2019). fMRIPrep: A robust preprocessing pipeline for functional MRI. Nature Methods, 16(1), 111–116. 10.1038/s41592-018-0235-4

Gläscher, J., Daw, N., Dayan, P., & O’Doherty, J. P. (2010). States versus rewards: Dissociable neural prediction error signals underlying model-based and model-free reinforcement learning. Neuron, 66(4), 585–595. 10.1016/j.neuron.2010.04.016

Graybiel, A. M., & Grafton, S. T. (2015). The striatum: Where skills and habits meet. Cold Spring Harbor Perspectives in Biology, 7(8), 1–14. 10.1101/cshperspect.a021691

Guell, X., Gabrieli, J. D. E., & Schmahmann, J. D. (2018). Embodied cognition and the cerebellum: Perspectives from the Dysmetria of Thought and the Universal Cerebellar Transform theories. Cortex, 100, 140–148. 10.1016/j.cortex.2017.07.005

Heleven, E., van Dun, K., De Witte, S., Baeken, C., & Van Overwalle, F. (2021). The Role of the Cerebellum in Social and Non-Social Action Sequences: A Preliminary LF-rTMS Study. Frontiers in Human Neuroscience, 15. 10.3389/fnhum.2021.593821

Hoffman, L. J., Foley, J. M., Leong, J. K., Sullivan-Toole, H., Elliott, B. L., & Olson, I. R. (2024). A Virtual In Vivo Dissection and Analysis of Socioaffective Symptoms Related to Cerebellum-Midbrain Reward Circuitry in Humans. Journal of Neuroscience, 44(41). 10.1523/JNEUROSCI.1031-24.2024

Ito, M. (2008). Control of mental activities by internal models in the cerebellum. Nature Reviews Neuroscience, 9(4), 304–313. 10.1038/nrn2332

Jack, A., Keifer, C. M., & Pelphrey, K. A. (2017). Cerebellar contributions to biological motion perception in autism and typical development. Human Brain Mapping, 38(4), 1914–1932. 10.1002/hbm.23493

Jack, A., & Pelphrey, K. A. (2015). Neural Correlates of Animacy Attribution Include Neocerebellum in Healthy Adults. Cerebral Cortex, 25(11), 4240–4247. 10.1093/cercor/bhu146

Jarcho, J. M., Romer, A. L., Shechner, T., Galvan, A., Guyer, A. E., Leibenluft, E., Pine, D. S., & Nelson, E. E. (2015). Forgetting the best when predicting the worst: Preliminary observations on neural circuit function in adolescent social anxiety. Developmental Cognitive Neuroscience, 13, 21–31. 10.1016/j.dcn.2015.03.002

Kawato, M., Furukawa, K., & Suzuki, R. (1987). A hierarchical neural-network model for control and learning of voluntary movement. Biological Cybernetics, 57, 169–185.

King, M., Hernandez-Castillo, C. R., Poldrack, R. A., Ivry, R., & Diedrichsen, J. (2019). Functional boundaries in the human cerebellum revealed by a multi-domain task battery. Nature Neuroscience, 22(August). 10.1038/s41593-019-0436-x

Kostadinov, D., Beau, M., Blanco-Pozo, M., & Häusser, M. (2019). Predictive and reactive reward signals conveyed by climbing fiber inputs to cerebellar Purkinje cells. Nature Neuroscience, 22(6), 950–962. 10.1038/s41593-019-0381-8

Kruppa, J. A., Gossen, A., Oberwelland Weiß, E., Kohls, G., Großheinrich, N., Cholemkery, H., Freitag, C. M., Karges, W., Wölfle, E., Sinzig, J., Fink, G. R., Herpertz-Dahlmann, B., Konrad, K., & Schulte-Rüther, M. (2019). Neural modulation of social reinforcement learning by intranasal oxytocin in male adults with high-functioning autism spectrum disorder: A randomized trial. Neuropsychopharmacology, 44(4), 749–756. 10.1038/s41386-018-0258-7

Limperopoulos, C., Bassan, H., Gauvreau, K., Robertson, R. L., Sullivan, N. R., Benson, C. B., Avery, L., Stewart, J., Soul, J. S., Ringer, S. A., Volpe, J. J., & Du Plessis, A. J. (2007). Does cerebellar injury in premature infants contribute to the high prevalence of long-term cognitive, learning, and behavioral disability in survivors? Pediatrics, 120(3), 584–593. 10.1542/peds.2007-1041

Limperopoulos, C., Chilingaryan, G., Sullivan, N., Guizard, N., Robertson, R. L., & Du Plessis, A. J. (2014). Injury to the premature cerebellum: Outcome is related to remote cortical development. Cerebral Cortex, 24(3), 728–736. 10.1093/cercor/bhs354

Lin, A., Rangel, A., & Adolphs, R. (2012). Impaired Learning of Social Compared to Monetary Rewards in Autism. Frontiers in Neuroscience, 6. 10.3389/fnins.2012.00143

Manto, M., Adamaszek, M., Apps, R., Carlson, E., Guarque-Chabrera, J., Heleven, E., Kakei, S., Khodakhah, K., Kuo, S.-H., Lin, C.-Y. R., Joshua, M., Miquel, M., Mitoma, H., Larry, N., Péron, J. A., Pickford, J., Schutter, D. J. L. G., Singh, M. K., Tan, T., … Yamashiro, K. (2024). Consensus Paper: Cerebellum and Reward. The Cerebellum. 10.1007/s12311-024-01702-0

Metoki, A., Wang, Y., & Olson, I. R. (2022). The Social Cerebellum: A Large-Scale Investigation of Functional and Structural Specificity and Connectivity. Cerebral Cortex, 32(5), 987–1003. 10.1093/cercor/bhab260

Niv, Y. (2009). Reinforcement learning in the brain. Journal of Mathematical Psychology, 53(3), 139–154. 10.1016/j.jmp.2008.12.005

Olson, I. R., Hoffman, L. J., Jobson, K. R., Popal, H. S., & Wang, Y. (2023). Little brain, little minds: The big role of the cerebellum in social development. Developmental Cognitive Neuroscience, 60, 101238. 10.1016/j.dcn.2023.101238

Peak, J., Hart, G., & Balleine, B. W. (2019). From learning to action: The integration of dorsal striatal input and output pathways in instrumental conditioning. European Journal of Neuroscience, 49(5), 658–671. 10.1111/ejn.13964

Quarmley, M. E., Nelson, B. D., Clarkson, T., White, L. K., & Jarcho, J. M. (2019). I Knew You Weren’t Going to Like Me! Neural Response to Accurately Predicting Rejection Is Associated With Anxiety and Depression. Frontiers in Behavioral Neuroscience, 13. https://www.frontiersin.org/articles/10.3389/fnbeh.2019.00219

Rapee, R. M., & Heimberg, R. G. (1997). A cognitive-behavioral model of anxiety in social phobia. Behaviour Research and Therapy, 35(8), 741–756. 10.1016/S0005-7967(97)00022-3

Raymond, J. L., Lisberger, S. G., & Mauk, M. D. (1996). The cerebellum: A neuronal learning machine? Basic cerebellar circuitry. Science.

Raymond, J. L., & Medina, J. F. (2018). Computational principles of supervised learning in the cerebellum. Annual Review of Neuroscience, 41(1), 233–253. 10.1146/annurev-neuro-080317-061948

Richards, J. M., Plate, R. C., & Ernst, M. (2013). A systematic review of fMRI reward paradigms used in studies of adolescents vs. adults: The impact of task design and implications for understanding neurodevelopment. Neuroscience & Biobehavioral Reviews, 37(5), 976–991. 10.1016/j.neubiorev.2013.03.004

Rosenblau, G., Korn, C. W., & Pelphrey, K. A. (2018). A computational account of optimizing social predictions reveals that adolescents are conservative learners in social contexts. Journal of Neuroscience, 38(4), 974–988. 10.1523/JNEUROSCI.1044-17.2017

Sathyanesan, A., Zhou, J., Scafidi, J., Heck, D. H., Sillitoe, R. V., & Gallo, V. (2019). Emerging connections between cerebellar development, behaviour and complex brain disorders. Nature Reviews Neuroscience, 20(5), 298–313. 10.1038/s41583-019-0152-2

Schmahmann, J. D. (2019). The cerebellum and cognition. Neuroscience Letters, 688(April 2018), 62–75. 10.1016/j.neulet.2018.07.005

Schultz, W. (2000). Multiple reward signals in the brain. Nature Reviews Neuroscience, 1(3), 199–207. 10.1038/35044563

Shen, X., Helion, C., Smith, D. V., & Murty, V. P. (2024). Motivation as a Lens for Understanding Information-seeking Behaviors. Journal of Cognitive Neuroscience, 36(2), 362–376. 10.1162/jocn_a_02083

Smith, D. V., Clithero, J. A., Boltuck, S. E., & Huettel, S. A. (2014). Functional connectivity with ventromedial prefrontal cortex reflects subjective value for social rewards. Social Cognitive and Affective Neuroscience, 9(12), 2017–2025. 10.1093/scan/nsu005

Smith, D. V., Hayden, B. Y., Truong, T.-K., Song, A. W., Platt, M. L., & Huettel, S. A. (2010). Distinct Value Signals in Anterior and Posterior Ventromedial Prefrontal Cortex. Journal of Neuroscience, 30(7), 2490–2495. 10.1523/JNEUROSCI.3319-09.2010

Sokolov, A. A. (2018). The Cerebellum in Social Cognition. Frontiers in Cellular Neuroscience, 12. 10.3389/fncel.2018.00145

Supekar, K., Musen, M., & Menon, V. (2009). Development of Large-Scale Functional Brain Networks in Children. PLOS Biology, 7(7), e1000157. 10.1371/journal.pbio.1000157

Tricomi, E., & Fiez, J. A. (2012). Information content and reward processing in the human striatum during performance of a declarative memory task. *Cognitive, Affective*, & Behavioral Neuroscience, 12(2), 361–372. 10.3758/s13415-011-0077-3

Van Overwalle, F., D’aes, T., & Mariën, P. (2015). Social cognition and the cerebellum: A meta-analytic connectivity analysis. Human Brain Mapping, 36(12), 5137–5154. 10.1002/hbm.23002

Van Overwalle, F., & Mariën, P. (2016). Functional connectivity between the cerebrum and cerebellum in social cognition: A multi-study analysis. NeuroImage, 124, 248–255. 10.1016/j.neuroimage.2015.09.001

Van Overwalle, F., Van de Steen, F., & Mariën, P. (2019). Dynamic causal modeling of the effective connectivity between the cerebrum and cerebellum in social mentalizing across five studies. *Cognitive*, Affective and Behavioral Neuroscience, 19(1), 211–223. 10.3758/s13415-018-00659-y

Van Overwalle, F., Van de Steen, F., van Dun, K., & Heleven, E. (2020). Connectivity between the cerebrum and cerebellum during social and non-social sequencing using dynamic causal modelling. NeuroImage, 206(July 2019), 116326. 10.1016/j.neuroimage.2019.116326

Wagner, M. J., Kim, T. H., Savall, J., Schnitzer, M. J., & Luo, L. (2017). Cerebellar granule cells encode the expectation of reward. Nature, 544(7648), 96–100. 10.1038/nature21726

Wang, S. S. H., Kloth, A. D., & Badura, A. (2014). The cerebellum, sensitive periods, and autism. Neuron, 83(3), 518–532. 10.1016/j.neuron.2014.07.016

Wise, R. A. (2004). Dopamine, learning and motivation. Nature Reviews Neuroscience, 5(6), 483–494. 10.1038/nrn1406

Wolpert, D. M., & Ghahramani, Z. (2000). Computational principles of movement neuroscience. Nature Neuroscience, 3(11), 1212–1217. 10.1038/81497

Wolpert, D. M., Miall, R. C., & Kawato, M. (1998). Internal models in the cerebellum. Trends in Cognitive Sciences, 2(9), 338–347. 10.1007/s11116-016-9675-9

Wyngaarden, J. B., Johnston, C. R., Sazhin, D., Dennison, J. B., Zaff, O., Fareri, D., McCloskey, M., Alloy, L. B., Smith, D. V., & Jarcho, J. M. (2024). Corticostriatal responses to social reward are linked to trait reward sensitivity and subclinical substance use in young adults. Social Cognitive and Affective Neuroscience, 19(1), nsae033. 10.1093/scan/nsae033

Yarkoni, T., Poldrack, R. A., Nichols, T. E., Van Essen, D. C., & Wager, T. D. (2011). Large-scale automated synthesis of human functional neuroimaging data. Nature Methods, 8(8), 665–670. 10.1038/nmeth.1635

