## Supplementary materials for "The Cerebellum and Striatum in Reward Processing: Caring About Being Right vs. Caring About Reward"

### METHODS

#### MRI Preprocessing

Results included in this manuscript come from preprocessing performed using *fMRIPrep* 22.0.2 (Esteban, Blair, et al., 2019; Esteban, Markiewicz, et al., 2019) which is based on *Nipype* 1.8.5 (Gorgolewski et al., 2011).

##### Anatomical data preprocessing

A total of 1 T1-weighted (T1w) images were found within the input BIDS dataset. The T1-weighted (T1w) image was corrected for intensity non-uniformity (INU) with N4BiasFieldCorrection (Tustison et al. 2010), distributed with ANTs 2.3.3 (Avants et al. 2008, RRID:SCR_004757), and used as T1w-reference throughout the workflow. The T1w-reference was then skull-stripped with a *Nipype* implementation of the antsBrainExtraction.sh workflow (from ANTs), using OASIS30ANTs as target template. Brain tissue segmentation of cerebrospinal fluid (CSF), white-matter (WM) and gray-matter (GM) was performed on the brain-extracted T1w using fast (FSL 6.0.5.1:57b01774, RRID:SCR_002823, Zhang, Brady, and Smith 2001). Brain surfaces were reconstructed using recon-all (FreeSurfer 7.2.0, RRID:SCR_001847, Dale, Fischl, and Sereno 1999), and the brain mask estimated previously was refined with a custom variation of the method to reconcile ANTs-derived and FreeSurfer-derived segmentations of the cortical gray-matter of Mindboggle (RRID:SCR_002438, Klein et al. 2017). Volume-based spatial normalization to one standard space (MNI152NLin2009cAsym) was performed through nonlinear registration with antsRegistration (ANTs 2.3.3), using brain-extracted versions of both T1w reference and the T1w template. The following template was selected for spatial normalization: *ICBM 152*

*Nonlinear Asymmetrical template version 2009c* [Fonov et al. (2009), RRID:SCR_008796; TemplateFlow ID: MNI152NLin2009cAsym].

#### Functional data preprocessing

For each of the 4 BOLD runs found per subject (across all tasks and sessions), the following preprocessing was performed. First, a reference volume and its skull-stripped version were generated using a custom methodology of *fMRIPrep*. Head-motion parameters with respect to the BOLD reference (transformation matrices, and six corresponding rotation and translation parameters) are estimated before any spatiotemporal filtering using mcflirt (FSL 6.0.5.1:57b01774, Jenkinson et al. 2002). BOLD runs were slice-time corrected to 1.01s (0.5 of slice acquisition range 0s-2.02s) using 3dTshift from AFNI (Cox and Hyde 1997, RRID:SCR_005927). The BOLD time-series (including slice-timing correction when applied) were resampled onto their original, native space by applying the transforms to correct for head-motion. These resampled BOLD time-series will be referred to as *preprocessed BOLD in original space*, or just *preprocessed BOLD*. The BOLD reference was then co-registered to the T1w reference using bbregister (FreeSurfer) which implements boundary-based registration (Greve and Fischl 2009). Co-registration was configured with six degrees of freedom. Several confounding time-series were calculated based on the *preprocessed BOLD*: framewise displacement (FD), DVARS and three region-wise global signals. FD was computed using two formulations following Power (absolute sum of relative motions, Power et al. (2014)) and Jenkinson (relative root mean square displacement between affines, Jenkinson et al. (2002)). FD and DVARS are calculated for each functional run, both using their implementations in *Nipype* (following the definitions by Power et al. 2014). The three global signals are extracted within the CSF, the WM, and the whole-brain masks. Additionally, a set of physiological regressors were extracted to allow for component-based noise correction (*CompCor*, Behzadi et al. 2007). Principal components are estimated after high-pass filtering the *preprocessed BOLD* time-series (using a discrete cosine filter with 128s cut-off) for the two *CompCor* variants: temporal (tCompCor) and anatomical (aCompCor). tCompCor components are then calculated from the top 2% variable voxels within the brain mask. For aCompCor, three probabilistic masks (CSF, WM and combined CSF+WM) are generated in anatomical space. The implementation differs from that of Behzadi et al. in that instead of eroding the masks by 2 pixels on BOLD space, a mask of pixels that likely contain a volume fraction of GM is subtracted from the aCompCor masks. This mask is obtained by dilating a GM mask extracted from the FreeSurfer's *aseg* segmentation, and it ensures components are not extracted from voxels containing a minimal fraction of GM. Finally, these masks are resampled into BOLD space and binarized by thresholding at 0.99 (as in the original implementation). Components are also calculated separately within the WM and CSF masks. For each CompCor decomposition, the *k* components with the largest singular values are retained, such that the retained components' time series are sufficient to explain 50 percent of variance across the nuisance mask (CSF, WM, combined, or temporal). The remaining components are dropped from consideration. The head-motion estimates calculated in the correction step were also placed within the corresponding confounds file. The confound time series derived from head motion estimates and global signals were expanded with the inclusion of temporal derivatives and quadratic terms for each (Satterthwaite et al. 2013). Frames that exceeded a threshold of 0.5 mm FD or 1.5 standardized DVARS were annotated as motion outliers. Additional nuisance timeseries are calculated by means of principal components analysis of the signal found within a thin band (*crown*) of voxels around the edge of the brain, as proposed by (Patriat, Reynolds, and

Birn 2017). The BOLD time-series were resampled into standard space, generating a *preprocessed BOLD run in MNI152NLin2009cAsym space*. First, a reference volume and its skull-stripped version were generated using a custom methodology of *fMRIPrep*. All resamplings can be performed with *a single interpolation step* by composing all the pertinent transformations (i.e. head-motion transform matrices, susceptibility distortion correction when available, and co-registrations to anatomical and output spaces). Gridded (volumetric) resamplings were performed using antsApplyTransforms (ANTs), configured with Lanczos interpolation to minimize the smoothing effects of other kernels (Lanczos 1964). Non-gridded (surface) resamplings were performed using mri_vol2surf (FreeSurfer).

#### Psychophysiological Interaction Analysis

A psychophysiological interaction analysis was used to examine whether regions of the cerebellum were functionally correlated with regions of the cerebrum, during task conditions. For this analysis, a regressor for activity within an ROI of the cerebellum was created. Using a generalized PPI framework (McLaren et al., 2012), an interaction term was created between the cerebellum ROI activity and the all wins and all losses regressors, separately. After conducting the first-level GLM, a contrast was created between the two interaction terms. This resulted in a whole-brain map indicating when regions of the cerebrum were correlated with activity in the cerebellum, for when cerebellum activity was greater for wins versus losses, for each participant. Subject-level connectivity maps were averaged together, and a one-sample t-test was done for each voxel in the brain to test whether the correlation was significantly greater than 0. Finally, an FDR correction was done to account for multiple comparisons, using a family-wise error rate of $p < .05$.

#### Model-based Representational Similarity Analysis

A neural representational dissimilarity matrix (RDM) was created using multi-voxel patterns of activity, across an ROI. Activity from voxels within an ROI were extracted from t-statistic maps for each of the conditions and runs in each task. The responses for each condition were correlated to each other for a given ROI across runs, but not within the same run. Then, dissimilarity was calculated by subtracting the r-value between each pair of conditions from 1, resulting in a neural RDM for each ROI. Neural RDMs were then correlated to the hypothesis RDM, using Spearman's rho, to test whether an ROI processed positive wins similarly to negative wins. First, a hypothesis representational dissimilarity matrix (RDM) was created. This RDM is a six by six matrix which mathematically represents the hypothesized similarity between our conditions of interest (Fig. 3). To test the hypothesis that the cerebellum is a domain-general error processor, a zero was inputted into the hypothesis RDM for the positive win to negative win and the positive loss to negative loss comparisons. This hypothesizes that the a region would have a similar response to positive wins as negative wins, and positive losses to negative losses. All other condition-pair comparisons were 1, indicating that a brain region would have a different response for each of the other conditions.

### Results

#### Univariate Analysis

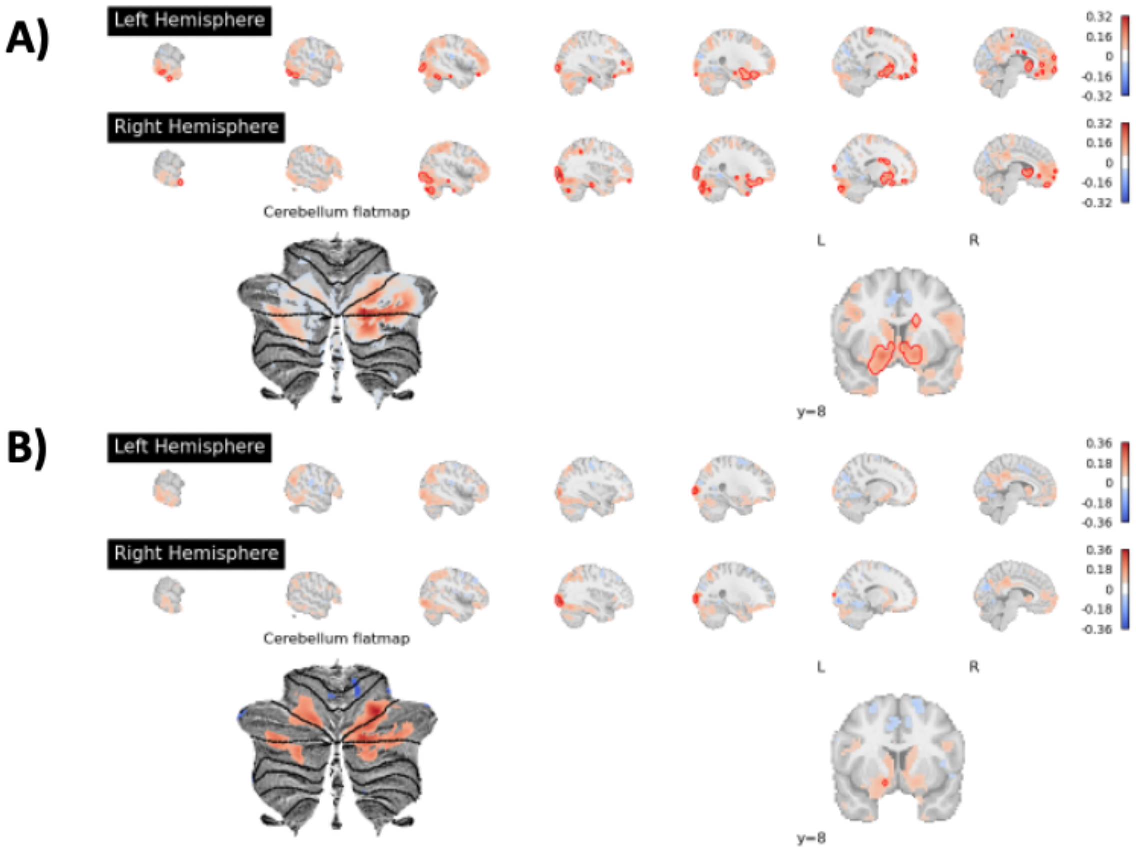


**Supplementary Figure 1.** Whole-brain univariate results for the monetary reward task, including all adolescent and young adult participants. A) Positive wins > positive losses contrast. B) All wins > all losses contrast. After accounting for multiple comparisons correction, using an FDR correction of $p < .05$, significant results are outlined in solid red lines, while the uncorrected results are shown as transparent, weighted by effect size.

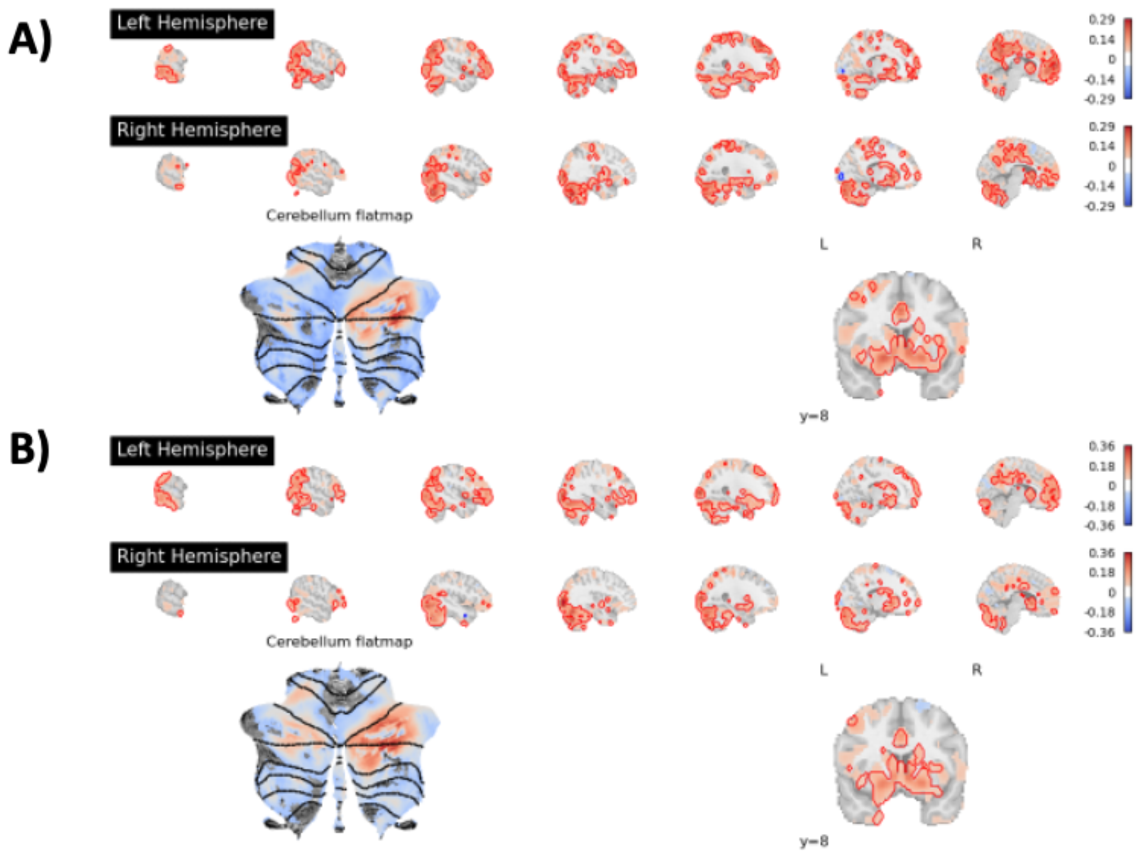


**Supplementary Figure 2.** Whole-brain univariate results for the social reward task, including all adolescent and young adult participants. A) Positive wins > positive losses contrast. B) All wins > all losses contrast. After accounting for multiple comparisons correction, using an FDR correction of $p < .05$, significant results are outlined in solid red lines, while the uncorrected results are shown as transparent, weighted by effect size.

#### Psychophysiological Interaction Analysis

Two regions within the cerebellum, CB region 8 and 9, were shown to be sensitive to wins vs losses in both the positive and all contrasts of the social reward task (Fig. 4). The CB region 8 ROI was selected as the ROI for the PPI analysis since it had slightly higher, although not significant, activity in both contrasts. The between group PPI analysis did not reveal any significant differences between young adults and adolescents (Supplementary Fig. 3). Within each group, there were no regions which survived multiple comparisons correction. However, interesting patterns of connectivity can be seen in the uncorrected maps. The young adults seem to have negative connectivity between CB region 8 and regions of the ventral visual stream in the social reward task (Supplementary Fig. 5A), while the adolescents show positive connectivity with regions of the social brain network including the PCC, vmPFC, and ATL (Supplementary Fig. 5B).

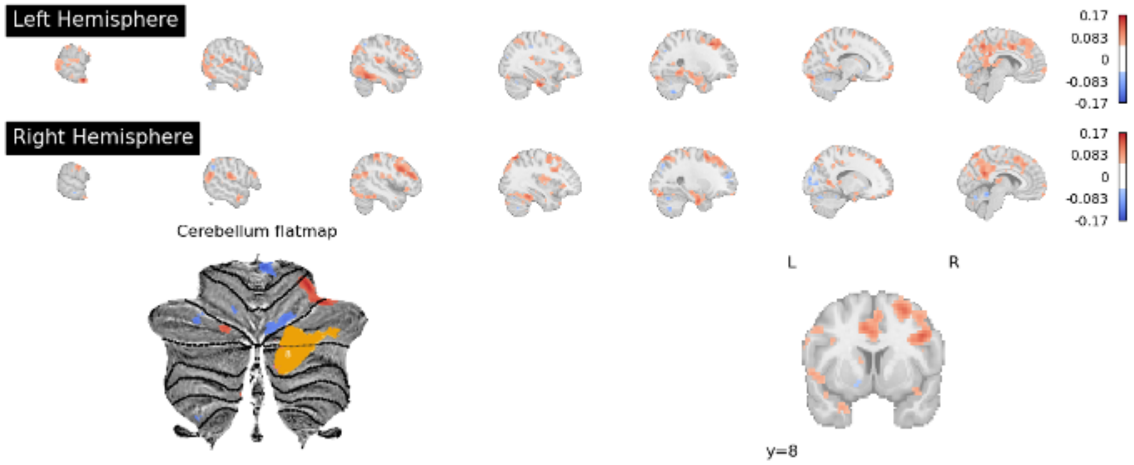


**Supplementary Figure 3.** Between group psychophysiological interaction analysis results for the social reward task, with red indicating greater connectivity for adolescents than young adults. Region 8 from the multi-domain task battery cerebellum atlas was used as an ROI, shown in yellow in the cerebellum flatmap. Maps are uncorrected for multiple comparisons.

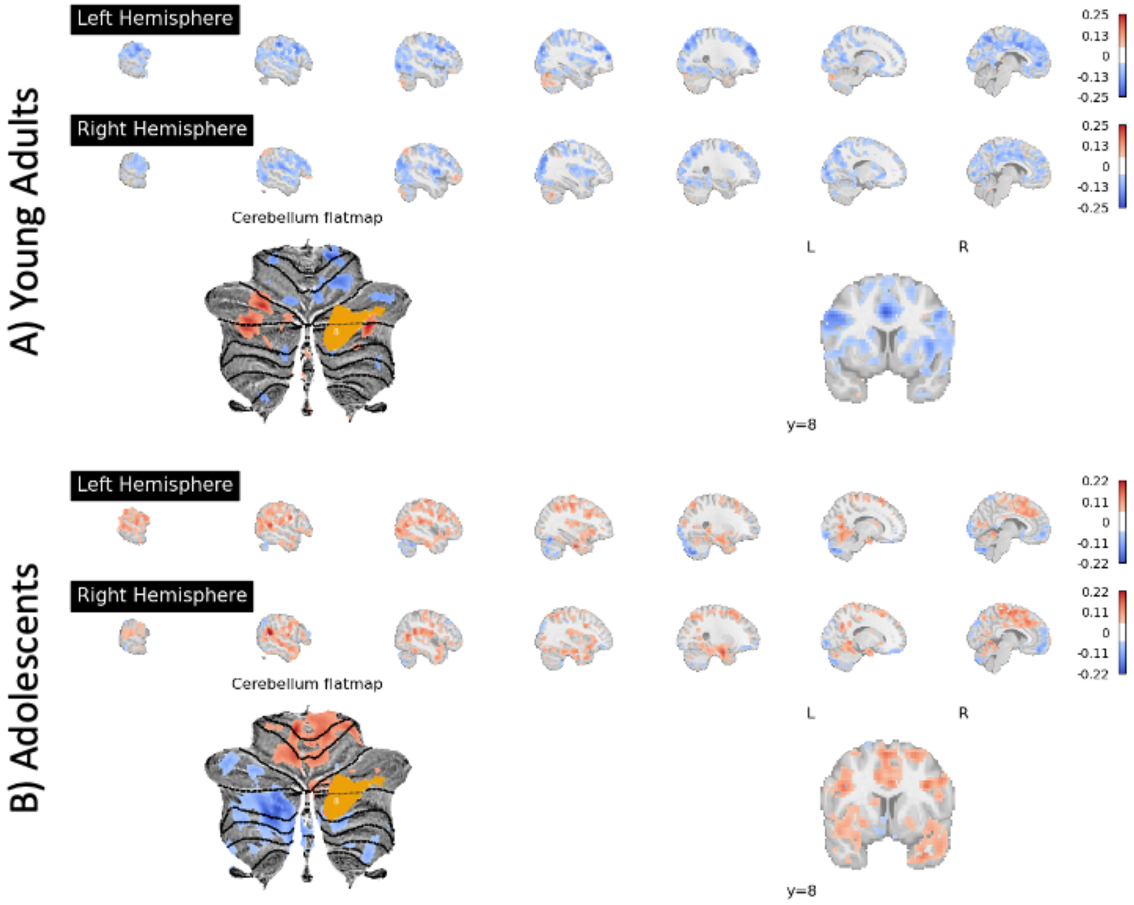


**Supplementary Figure 4.** Psychophysiological interaction analysis results for the monetary reward task in A) young adults, and B) adolescents. Region 8 from the multi-domain task battery cerebellum atlas was used as an ROI, shown in yellow in the cerebellum flatmap. Maps are uncorrected for multiple comparisons.

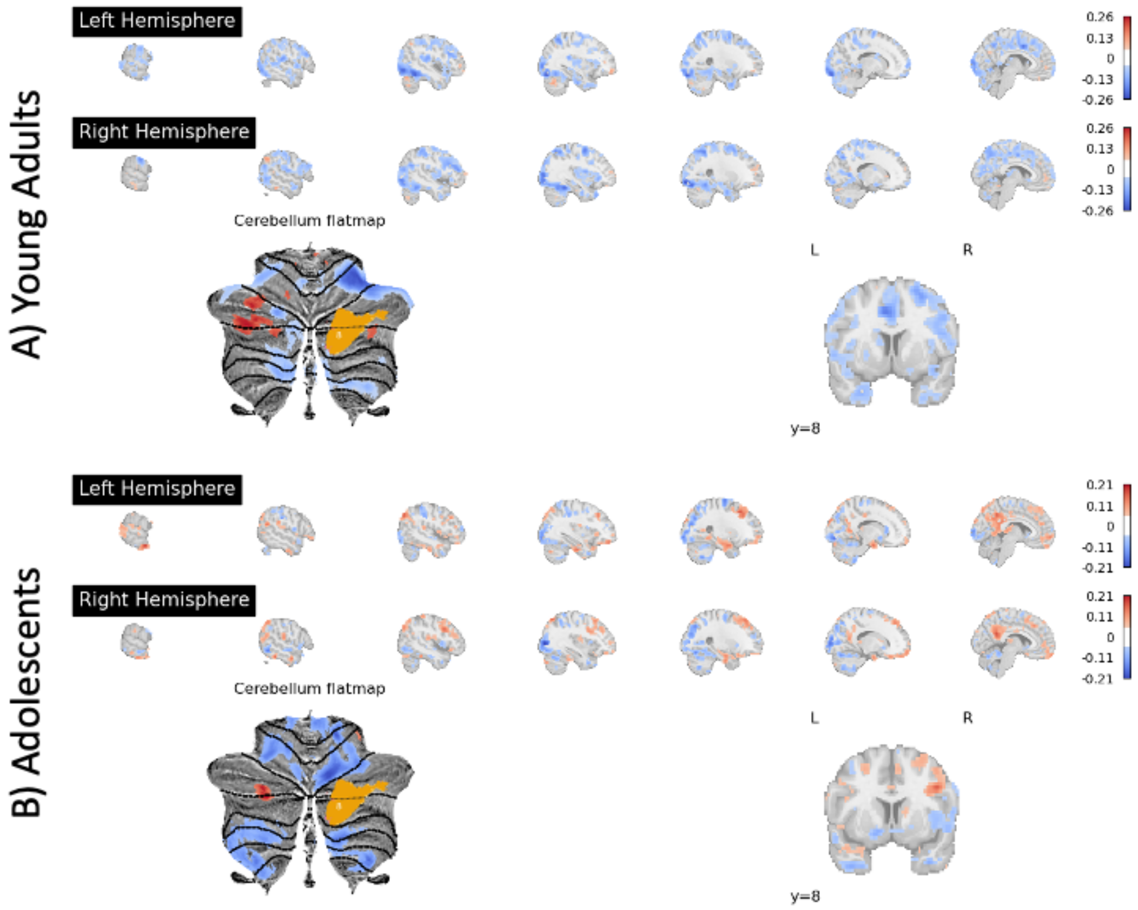


**Supplementary Figure 5.** Psychophysiological interaction analysis results for the social reward task in A) young adults, and B) adolescents. Region 8 from the multi-domain task battery cerebellum atlas was used as an ROI, shown in yellow in the cerebellum flatmap. Maps are uncorrected for multiple comparisons.

#### Representational Similarity Analysis

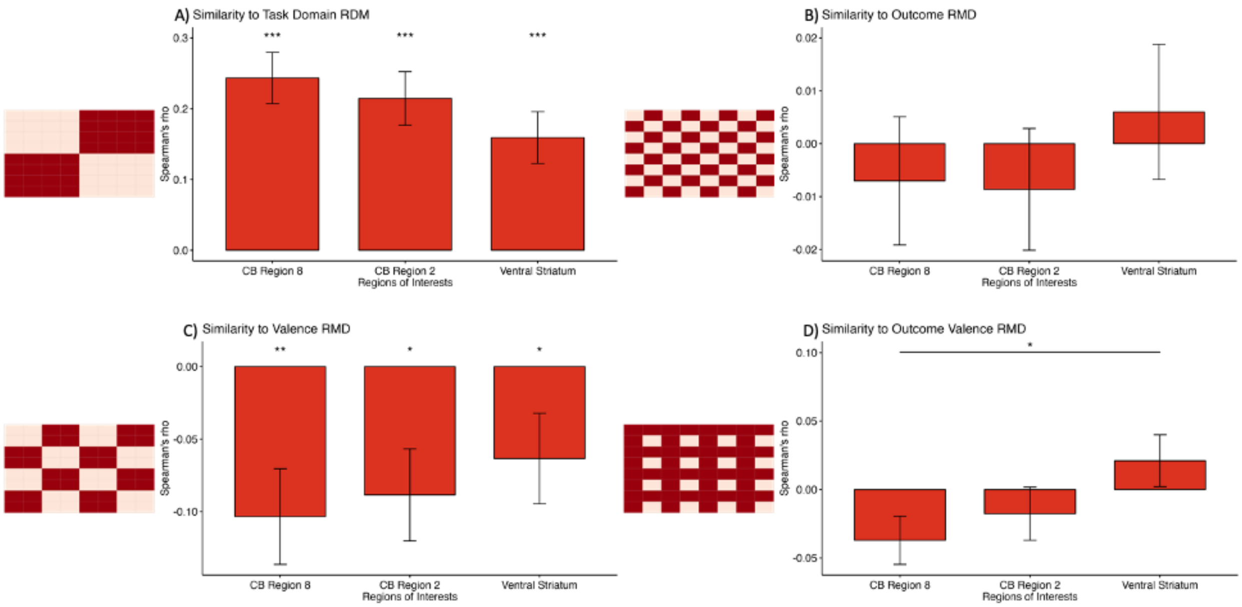


**Supplementary Figure 6.** Model representational similarity analysis. A) Correlation to task domain model RDM, hypothesizing greater similarity for feedback conditions within the same task (e.g. r(social, social) > r(social, monetary)). B) Correlation to outcome model RDM, hypothesizing greater similarity for feedback of the same outcome (e.g. r(win, win) > r(win, loss)). C) Correlation to valence model RDM, hypothesizing greater similarity for feedback of the same valence (e.g. r(positive, positive) > r(positive, negative)). D) Correlation to outcome valence model RDM, hypothesizing only similarity for positive loss and negative loss.

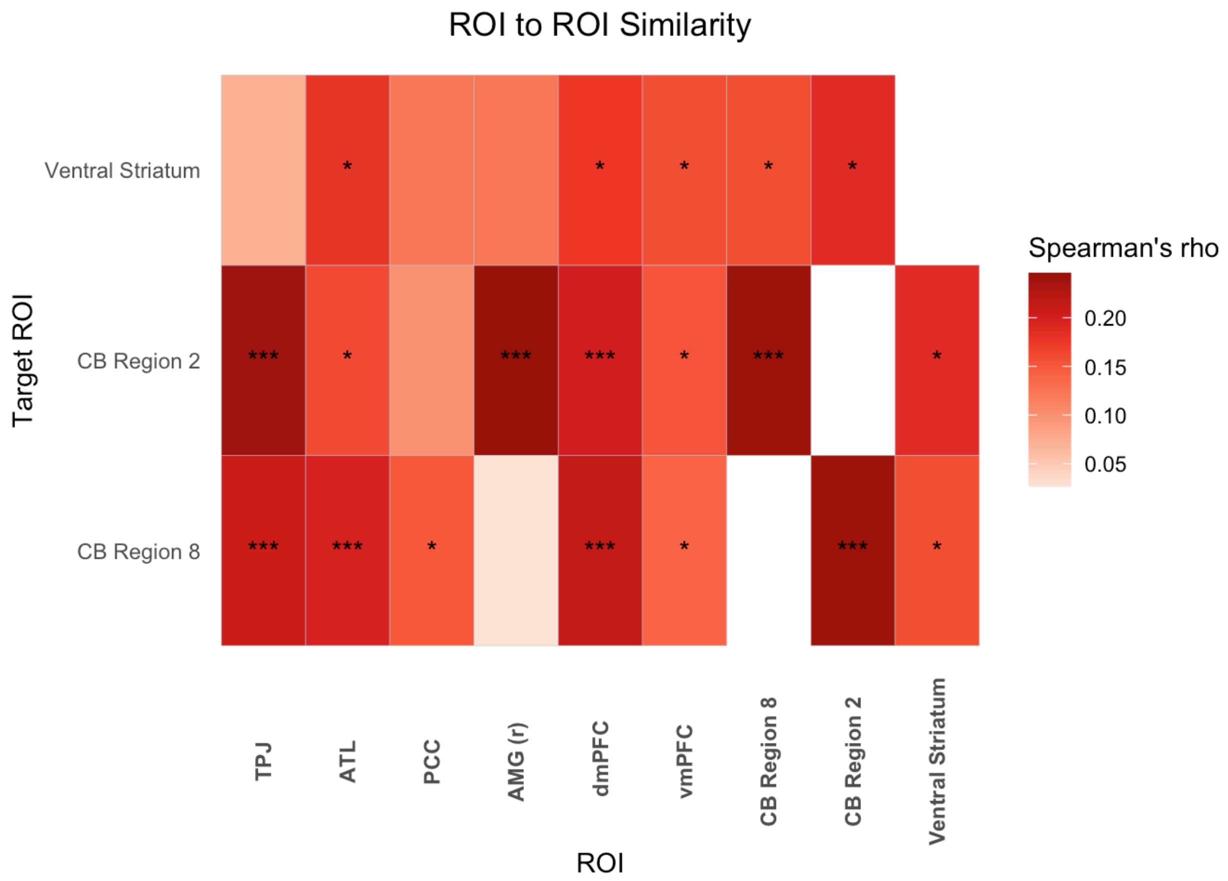


**Supplementary Figure 7.** ROI to ROI RSA. The neural RDMs for three target ROIs (ventral striatum, CB region 2, and CB region 8) were correlated to each other and ROIs of the social network.
